# A complex acoustical environment during development enhances auditory perception and coding efficiency in the zebra finch

**DOI:** 10.1101/2024.06.25.600670

**Authors:** Samantha M Moseley, C Daniel Meliza

## Abstract

Sensory experience during development has lasting effects on perception and neural processing. Exposing juvenile animals to artificial stimuli influences the tuning and functional organization of the auditory cortex, but less is known about how the rich acoustical environments experienced by vocal communicators affect the processing of complex vocalizations. Here, we show that in zebra finches (*Taeniopygia guttata*), a colonial-breeding songbird species, exposure to a naturalistic social-acoustical environment during development has a profound impact on auditory perceptual behavior and on cortical-level auditory responses to conspecific song. Compared to birds raised by pairs in acoustic isolation, male and female birds raised in a breeding colony were better in an operant discrimination task at recognizing conspecific songs with and without masking colony noise. Neurons in colony-reared birds had higher average firing rates, selectivity, and discriminability, especially in the narrow-spiking, putatively inhibitory neurons of a higher-order auditory area, the caudomedial nidopallium (NCM). Neurons in colony-reared birds were also less correlated in their tuning and more efficient at encoding the spectrotemporal structure of conspecific song, and better at filtering out masking noise. These results suggest that the auditory cortex adapts to noisy, complex acoustical environments by strengthening inhibitory circuitry, functionally decoupling excitatory neurons while maintaining overall excitatory-inhibitory balance.

**Significance Statement:** The statistics of the sensory inputs animals experience during postnatal development shape cortical circuits and their functional properties, but most studies examining experience-dependent plasticity in the auditory system has employed artificial stimuli with limited relevance to acoustic communication. Here, we examined how the natural social-acoustical environment experienced by zebra finches, a social songbird that breeds in large colonies, influences the development of auditory perception and the underlying auditory cortical circuits. Compared to birds raised in a more impoverished environment, colony-reared birds were better at recognizing songs of other zebra finches and had higher firing rates in the avian homolog to auditory cortex, along with changes to functional connectivity that resulted in more efficient and robust coding of conspecific song.

## Introduction

Experience is critical to the development of auditory perception. Human infants learn phonetic categories in the first year of life by hearing speech, resulting in changes to perception (Werker and Lalonde, 1988; Kuhl et al., 1992) paralleled by neural responses to speech sounds (Bidelman et al., 2013; Bosseler et al., 2013). In rodents, auditory experience during a developmental critical period has long-lasting effects on the functional organization of the auditory cortex (Zhang et al., 2001; de Villers-Sidani et al., 2007; Zhou and Merzenich, 2008; Insanally et al., 2009; Bao et al., 2013). In most songbird species, auditory experience not only provides a model to copy (Marler and Tamura, 1964; Gobes et al., 2019) but shapes how the auditory system processes conspecific vocalizations (Sturdy et al., 2001; Amin et al., 2013; Chen et al., 2017; Woolley, 2012). Because auditory perception is the foundation for vocal production and for higher-order aspects of communication, it is important to understand how natural experience shapes its development.

For vocal communicators that raise their young in groups, the sounds produced by conspecific individuals present opportunities and challenges. An enriched sensory environment is broadly beneficial to neuronal development (Woolley, 2012), and hearing many different individuals may be essential for learning the statistical invariants of song or speech (Maye et al., 2002). However, when multiple individuals vocalize simultaneously, it creates an acoustical background with identical statistics to the signals of interest (Cherry, 1953), which is a particularly challenging source of interference that the developing brain has to learn to filter out.

In many cases, the auditory system adapts to the environment in a manner consistent with Hebbian plasticity, which predicts that sensory neurons will become tuned to frequently experienced acoustical features or combinations of features. In zebra finches (*Taeniopygia guttata*) fostered by other finch species, cortical-level neurons are preferentially tuned to the spectrotemporal features of the foster species’ song (Moore and Woolley, 2019). In other cases, tuning shifts away from prevalent features of the acoustical environment, such as when rats are reared in spectrotemporally modulated noise (Homma et al., 2020). This anti-Hebbian plasticity may support the ability of animals to hear and discriminate vocalizations in natural environments, which are characterized by broadband, acoustically complex sounds from other animals and non-biological sources that can mask signals of interest (Waser and Brown, 1986; Brumm and Slabbekoorn, 2005), but the underlying mechanisms remain poorly understood.

In this study, we examined how a complex acoustical environment affects the development of auditory processing in the zebra finch, a social songbird that raises its young in dense colonies of tens to hundreds of individuals (Zann, 1996). Zebra finches communicate vocally using calls and songs with rich spectrotemporal structure (Elie and Theunissen, 2016) and are able to recognize other individuals by their songs and calls (Elie and Theunissen, 2018). Cortical-level auditory areas play a critical role in decoding individual identity and other messages from these signals (Wang et al., 2007; Schneider and Woolley, 2013; Elie and Theunissen, 2015). Raising zebra finches in continuous white noise results in abnormal cortical-level auditory responses (Amin et al., 2013), and exposing adults to the acoustical background of a canary colony transiently affects lateralization and frequency tuning (Yang and Vicario, 2015). However, the lasting impacts of the social-acoustical environment zebra finches typically experience throughout development remain unknown. Here, we compared birds raised in an indoor breeding colony (colony-reared, CR) to birds raised in acoustic isolation (pair-reared, PR), examining behavioral discrimination abilities and cortical-level auditory responses (in separate groups of animals). The CR birds were better at recognizing songs of other individuals, even when they were embedded in colony noise, and their neurons had higher firing rates, selectivity, and discriminability, especially the narrow-spiking, putatively inhibitory neurons of the caudomedial nidopallium (NCM). There were also effects on the relationship between signal and noise correlations that suggest exposure to a complex acoustical environment promotes development of inhibitory circuitry that decorrelates auditory responses, thereby improving the encoding of songs masked by colony noise.

## Materials and Methods

### Animals

All procedures were performed according to National Institutes of Health guidelines and protocols approved by the University of Virginia Institutional Animal Care and Use committee. Male and female zebra finches (*Taeniopygia guttata*) were bred in our local colony from 16 different pairs. All birds received finch seed (Abba Products, Hillside, NJ) and water ad libitum and were kept on a 16:8 light-dark schedule in temperature and humidity-controlled rooms (22–24 °C).

### Experimental Rearing Conditions

Zebra finches were reared in individual cages that were initially placed in the colony room, which housed around 70 male and female finches of varying ages at the time of the study. In the colony-reared (CR) condition, families remained in the colony room. In the pair-reared (PR) condition, families were moved to an acoustic isolation chamber (Eckel Industries) five days after the first chick hatched. Hatchlings were fed by hand after the move as needed. In both conditions, the parents were separated from their chicks at 35 dph (days post hatch). CR juveniles continued to be housed in the colony with other juveniles or in large single-sex flight aviaries. PR chicks remained with their siblings in the acoustic isolation chamber until they were used in an experiment.

### Acoustic Recordings and Analysis

Sound recordings were taken of the acoustical environment for one CR clutch and one PR clutch. A small hole was drilled in the lid of the nest box, a piece of foam rubber padding with a cavity for a microphone was glued above the hole, and an omnidirectional lavalier microphone (Shure SM93) was placed in the cavity. The signal was amplified and digitized at 48 kHz using a Focusrite Scarlett 2i2 and recorded directly to disk using custom software (jrecord v2.1.5 or v2.1.9, https://github.com/melizalab/jill). Before installing the lid on the nestbox, a recording was made of a 1 kHz calibration tone emitted by an R8090 (Reed Instruments) calibrator, which was placed immediately above the hole on the nest side of the lid. The amplitude of this tone at the opening of the R8090 was 80 dB SPL as measured by a calibrated sound meter (NTi XL2 with an M4261 microphone). The gain on the amplifier/digitizer was kept constant throughout the recording. The CR clutch comprised four chicks and was recorded for 7 days starting when the chicks were 26–29 dph. The PR clutch comprised six chicks and was recorded for 33 days starting when the chicks were 4–11 dph; however, only the recordings made when the chicks were a comparable age to the CR birds were used in this analysis.

To convert the units of the recordings to dB SPL, the power spectrum of the calibration recording was computed using Welch’s method with a 10 ms flat-top window, and a scaling factor was determined for the 80 dB SPL peak at 1 kHz. The amplitude statistics of the nestbox recordings were analyzed in segments of 10 min that overlapped by 50%. For each segment, the calibration scaling factor was applied, followed by an A-weighting digital filter. The segments were further subdivided into 10 ms frames with 50% overlap, and a Fourier transform with a Hanning window was used to estimate the power spectral density (PSD). The PSD in each frame was integrated and log-transformed to obtain amplitude on the dB_A_ SPL scale. The 0% (minimum), 25%, 50%, 75%, and 100% (maximum) quantiles of the amplitude were then computed for each 10-min segment.

The acoustical structure of the colony noise was analyzed using sound texture statistics based on a model of auditory processing that separates acoustic signals into log-spaced frequency bands corresponding to locations on the basilar membrane, extracts the amplitude envelope in each band on a nonlinearly compressed scale, and then further analyzes each acoustic frequency band into channels corresponding to different modulation rates (McDermott and Simoncelli, 2011). Statistics for the colony were computed from sixty-six 10 s segments sampled from the CR nestbox recording. The recording was divided into hours, and one sample was chosen at a random time from each of 66 daylight hours. One sample was excluded because it included sounds of the animal care staff physically manipulating the cage, which was a relatively rare occurrence. For comparison, texture statistics were calculated for 62 samples of zebra finch song from 12 individual males (5 or 6 songs per bird), drawn from the lab’s collection of recordings. These recordings were obtained in acoustic isolation chambers using the same digitization software and hardware but a different microphone (AudioTechnica Pro70). Both the SM93 and the Pro70 have frequency response curves that are flat to within ±10 dB between 100–14,000 Hz. Song samples were chosen to be 10 s long; for some birds with shorter songs, these samples comprised bouts of 2–3 motifs separated by gaps of 1–2 seconds. For both colony noise and song, texture statistics were calculated separately for each segment and then averaged.

Texture statistics were calculated using the same algorithm and parameters reported by McDermott and Simoncelli (2011), except that our acoustic frequency channels spanned from 246– 9005 Hz, as the lower frequencies in our recordings of colony noise or individuals did not contain any vocal signals. We reimplemented the algorithm in Python so that we could efficiently perform analysis on the large number of samples used in this study (https://github.com/melizalab/colony-noise, to be released on acceptance of the manuscript). We confirmed that the output of our implementation matched that of the MATLAB code provided by McDermott and Simoncelli (http://mcdermottlab.mit.edu/downloads.html).

### Stimuli

The stimuli comprised songs selected from recordings of 10 male zebra finches. To minimize familiarity, we used birds from the Margoliash lab at the University of Chicago (*n* = 7) or birds in our colony that never reproduced (*n* = 3). Songs included 1–2 motifs and were 2.52 ± 0.28 s in duration (mean ± SD; range 1.92–2.88 s). The songs were filtered with a 4th-order Butterworth highpass filter with a cutoff of 150 Hz, a 2 ms squared cosine ramp was applied to the beginnings and the ends, and the waveforms were rescaled to have an RMS amplitude of –30 dB relative to the full scale of the sound file. The songs were then embedded in background synthetic noise generated by the McDermott and Simoncelli (2011) algorithm to match the statistics of our colony nestbox recordings. The amplitude of the background was varied to give signal-to-noise ratios (SNRs) ranging from 70 to –10 dB. The foreground songs were presented at approximately 60 dB_A_ SPL in both behavioral and electrophysiological experiments, so the noise at the highest SNR level (–10 dB_A_ SPL) was well below the hearing threshold of the finches (Hashino and Okanoya, 1989).

### Behavior

One cohort of birds was tested for their ability to discriminate conspecific songs embedded in synthetic colony noise. At 60–98 dph (mean ± SD: 75 ± 12 dph), birds were individually housed in acoustic isolation chambers, each containing an operant apparatus with a speaker (iML227 Orbit USB Lite, Altec Lansing) and a custom response panel with three keys and a food dispenser (https://meliza.org/starboard/). After 3 days to acclimate, the birds were shaped to initiate trials by pecking the center key and then to peck either the left or right key when it was lit to obtain a seed reward. After successfully completing 100 shaping trials, the birds were pre-trained to peck the left key in response to one zebra finch song and peck the right key in response to another song. Birds had 5 s to respond after the end of the stimulus. Pecks to the correct key were rewarded; pecks to the wrong key and failures to respond were punished by turning the lights in the box out for 10 s, during which time the bird could not initiate a new trial. To correct key biases, incorrect responses were followed by up to 50 correction trials in which the same stimulus was presented until the bird made the correct response. On some correction trials a light cueing the correct answer was added; this support was gradually withdrawn as performance improved. Following pre-training, birds were transferred to a new set of four unique songs, with two songs assigned to one key and two to the other. There were two non-overlapping sets of training songs (A and B). Two of the four CR birds and two of the six PR birds were trained on set A. The assignment of stimuli within each training set to left or right response keys was counterbalanced. The gain on the speaker was adjusted so that the foreground songs had an RMS amplitude of 58 dB_A_ SPL at the location where the bird perched to initiate trials. Each song was embedded in five different samples of synthetic colony background, which started 500 ms before the onset of the song and ended 1000 ms after the song ended. During training, the SNR was 70 dB. Birds were excluded if they failed to shape or failed to reach 80% accuracy on all stimuli within 3000 trials (excluding corrections) during pre-training or training. The task was difficult for birds to learn: out of an initial cohort of 17 CR and 16 PR birds, only 4 CR (2 male) and 6 PR (2 male) birds were included in the analysis.

After the birds achieved 80% or better performance in training, we prepared them to discriminate the songs in a noisy background by introducing stimuli in which the SNR was 35 dB and reducing the probability of reinforcement to 80%. Correction trials and light cues were not used in this pre-test phase or in the testing that followed. After birds successfully matched performance on the 70 and 35 dB SNR stimuli, we introduced test stimuli with SNRs ranging from 30 to –10 dB using a block design. In each block, we tested one SNR level; 40% of the trials were at 70 dB SNR, 40% at 35 dB SNR, and 20% were at the test SNR level. We collected at least 20 trials of the test stimuli for each foreground song and then moved to the next block. Birds started at the easiest level (30 dB SNR) and were moved to the next harder SNR in steps of –5 dB. All the birds were tested up to –10 dB SNR except one, which we stopped after –5 dB SNR for expedience. Responses to all the stimuli were reinforced throughout training and testing.

### Electrophysiology

#### Surgery

A second cohort of 9 CR (4 male) and 7 PR (5 male) birds was used to examine the effects of the acoustical environment on neuronal response properties. Three days before recordings, adult birds (90–110 dph) were anesthetized with 3.5% isoflurane (Southmedic). Feathers were removed from the top of the head and the bird was placed in a stereotaxic holder. The scalp was prepared with 7.5% povidone iodine (McKesson), Neosporin (Johnson & Johnson), and 2% Lidocaine (Henry Schein) prior to incision. The recording site was identified using stereotaxic coordinates relative to the Y-sinus. A metal pin was affixed to the skull rostral to the recording site with dental acrylic, and the first layer of skull over the recording site was removed with a scalpel. Subjects were allowed to recover between three to four days before recordings.

On the day of recording, the bird was given three intramuscular injections of 20% urethane (0.5 mL/g; Sigma-Aldrich), separated by 20 min. The bird was restrained in a padded 50 mL conical tube, and the head pin was attached to a stand in the recording chamber (Eckel Industries). For most birds, a craniotomy was performed adjacent to the recording site for ground wire placement; in other recordings the ground wire was placed in the well surrounding the recording site. The second layer of skull was removed over the recording site, the dura was thinned using an Ultra Needle (Electron Microscopy Sciences), and a well was formed with Kwik-Cast around the recording site to hold agarose and phosphate-buffered saline.

#### Stimulus Presentation

Acoustic stimuli were presented with custom Python software (https://github.com/melizalab/open-ephys-audio; v0.1.7) through a Focusrite 2i2 analog-to-digital converter, a Samson Servo 120a amplifier, and a Behringer Monitor Speaker 1C. The gain on the amplifier was adjusted so that the foreground songs had an RMS amplitude of 60 dB_A_ SPL at the position of the bird’s head. The songs were combined into sequences, each consisting of 10 permutations following a balanced Latin square design (i.e., each song occurred once in each of the 10 positions and was preceded by a different song in each permutation). The songs within each sequence were separated by gaps of 0.5 s, and the sequences were presented with a gap of 5 s. The sequences were embedded in a background consisting of synthetic colony noise that began 2 s before the first song and ended 2 s after the end of the last song. We only used one sample of background noise, because permuting the order of the foreground songs meant each song would be presented against 10 different segments of the background.

#### Data Acquisition

Neural activity was recorded using 128-channel two-shank silicon electrodes (128 AxN Sharp, Masmanidis Lab) connected to an Intan RHD 128-channel headstage. Data were collected at 30 kHz by the Open Ephys Acquisition Board connected to a computer running Open Ephys GUI software (v0.5.3.1). The electrode was coated with DiI (ThermoFisher Scientific) and inserted at a dorso-caudal to ventro-rostral angle that confined the pentrations to one of three groups of auditory areas: the thalamorecipient areas L2a and L2b; the superficial areas L1, CLM (caudolateral mesopallium), and CMM (caudomedial mesopallium); and the deep/secondary areas L3 and NCM (caudomedial nidopallium). Recordings were only included if at least 80% of the contacts on the electrode were in one of these areas. After contacting the brain surface, the probe was advanced at 1 µm/s. A separate collection of zebra finch songs was played during electrode advancement. Electrode movement was stopped once the local field potentials and spiking indicated reliable auditory responses across the whole probe. Once the electrode was in position, the well was filled with 2% agarose (Sigma-Aldrich) and phosphate-buffered saline. After responses to the full set of stimuli were recorded, we either advanced the electrode by at least 1.1 mm so that all the recording sites would be in a new region of the brain, withdrew the electrode and inserted it in a new location, or terminated the recording. Recordings were made from 1–3 different areas per animal (median 2).

#### Histology

Immediately after recording, animals were administered a lethal intramuscular injection of Euthasol (Vibrac) and perfused transcardially with 10 U/mL solution of sodium heparin in PBS (in mM: 10 478 Na2HPO4, 154 NaCl, pH 7.4) followed by 4% formaldehyde (in PBS). Brains were removed from the skull, postfixed overnight in 4% paraformaldehyde, blocked along the midline, and embedded in 3% agar (Bio-Rad). Sections were cut at 60 µm on a vibratome and mounted on slides. After drying for 30 min, the sections were coverslipped with Prolong Gold with DAPI (Thermo Fisher, catalog P36934; RRID:SCR_05961). The sections were imaged using epifluorescence with DAPI and Texas Red filter cubes to locate the DiI-labeled penetrations. Because we only painted the electrode once at the beginning of the experiment and made sure to move it 200 µm or more between penetrations, we were able to reconstruct the locations of multiple recording sites per animal.

#### Spike Sorting and Classification

Spikes were sorted offline with Kilosort 2.5 (Pachitariu et al., 2016). Clusters were automatically excluded if they had a contamination score higher than 10 or contained fewer than 100 spikes (corresponding to a rate of 0.029 Hz over the 58 min session), as it was not possible to assess quality with so few events. Clusters were further curated by visual inspection (phy 2.0; https://github.com/cortex-lab/phy) for spheroid distribution in the feature space, very low refractory period violations in the autocorrelogram, stable firing rate throughout the recording, and a typical average spike waveform. Individual spikes were excluded as artifacts if their amplitude was more than 6 times the amplitude of the average spike from that unit. To classify units as narrow-spiking (NS) or broad-spiking (BS), the spike waveforms for each unit were averaged after resampling to 150 kHz using sinc interpolation. Taking the time of the maximum negative deflection (trough) as 0, the spike width was defined as the time of the maximum positive deflection (peak) following the trough, and the peak/trough ratio was measured from the (absolute) magnitude of the peak and trough. A Gaussian mixture model with two groups was fit to these features, and units were classified as narrow-spiking if they were assigned to the group with the lower spike width and higher peak/trough ratio. Nearly identical results were obtained if separate classifiers were fit for each brain area.

#### Experimental Design and Statistical Analysis

The main independent variable in this study was rearing condition (CR or PR). Birds were chosen from CR and PR clutches at random while attempting to maintain a balance of males and females. Sex and family were not included as analysis variables due to the limited sample size. The experimenter was aware of the rearing condition of each animal.

In the neurophysiology experiments, we also compared brain areas (L2a/L2b, L1/CM, or L3/NCM) and spike types (BS or NS). We attempted to sample from every brain area in each animal, but as this was not always possible due to limited recording time, we biased our selection towards areas that were underrepresented at the time of the recording. Spike waveforms were not under our control; we included every well-isolated single unit. A fixed set of 10 songs was presented at every recording site. We employed many dependent measures, as described below. The number of birds, recording sites, and units for each area and condition in the electrophysiology experiments are given in Table 1. Data were analyzed using custom Python and R code (https://github.com/melizalab/cr_pr_adults, to be released on manuscript acceptance).

**Table 1.**
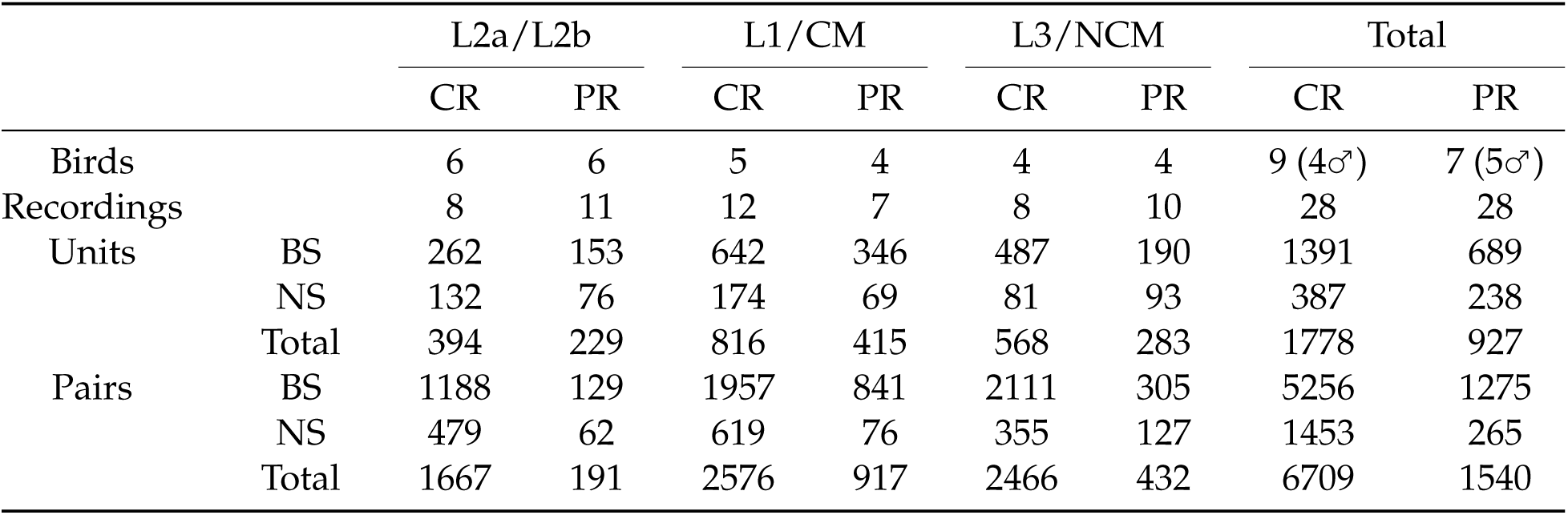
Sample sizes for birds, recordings, units, and pairs of simultaneously recorded units. Because only a subset of areas were sampled in most animals, the number of animals contributing data to each area varies. Each recording comprises responses to a complete stimulus set at a unique location in the auditory pallium. Units comprise all the well-isolated single units recorded in each area, and pairs comprise all the simultaneously recorded units in each area.

#### Behavior

The ability of birds to discriminate between stimuli during the training phase was computed using a state-space learning model (Smith et al., 2007). Briefly, the probability of pecking left on a given trial *t* was modeled as a Bernoulli random variable *y_t_* that depends on a latent state vector *x_t_* and the stimulus *Z_t_*:

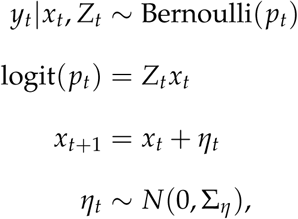

where *Z_t_* is a 2-element row vector that encodes the stimulus on trial *t*, with *Z_t_* = (1, −0.5) on trials where the stimulus is associated with the right key and (1, 0.5) on trials where the stimulus is associated with the left key. With this coding scheme, the state *x_t_* is a 2-element column vector representing the animal’s bias and its ability to discriminate left and right stimuli. Because of the logit transform, both components of *x_t_* are on the log-odds scale: bias is the log odds of the animal pecking the left key irrespective of the stimulus, and discrimination is the log odds ratio between pecking left for left-associated stimuli and right-associated stimuli. The state *x_t_* evolves as a multivariate Gaussian random walk with covariance Σ*_η_*. Given a series of *T* trials during the training phase, we can use Monte Carlo sampling to estimate the posterior distribution of Σ*_η_* and the sequence of latent states *x*_1:*T*_. We used the R package bssm (v2.0.2; Helske and Vihola, 2021) to generate 10,000 samples from the posterior latent state distributions for each behavioral subject. We used a separate state-space model for each subject to estimate how non-response probability changed over training. The state-space learning analysis was used for visualization only; because we made subject-specific changes to correction trial and light cue strategies to eliminate key biases and achieve a high level of discrimination in as many subjects as possible, a comparison of learning rate between groups is unlikely to be interpretable.

For the test trials, we estimated discrimination and non-response probability as a function of SNR with the assumption that there was no change in performance due to learning over the course of the experiment, which removed the need for a state-space model. To properly pool trials from different animals, we used a GLMM with a Bernoulli dependent variable, SNR and rearing condition as fixed effects, and a random intercept for each subject. SNR was encoded as a categorical variable, with 70 dB as the reference level. The parameters of the model were estimated in R using lme4 (v1.1-35). The model specification was `cbind(n_left, n_pecked - n_left) ∼ snr*rearing*stim_left + (1|subject)` for discrimination and `cbind(n_trials - n_pecked, n_trials) ∼ snr*rearing + (1|subject)` for non-response probability. Because the model is nonlinear, effects and confidence intervals are reported as estimated marginal means, calculated in R using the emmeans package (v1.10.1). Note that emmeans reports test statistics for post-hoc tests on GLMMs as z-scores, with infinite degrees of freedom. Contrasts are reported on the log odds ratio (LOR) scale.

#### Neurophysiology

Spike trains were separated into non-overlapping intervals corresponding to songs, beginning at the start of the song and ending 0.5 s after the end of the song. This gave us 10 trials per 10 songs at each of 11 SNR levels per unit. For spontaneous firing, we used 1 s intervals taken from the periods of silence at the beginning of each song sequence, for a total of 110 trials per unit.

Spontaneous and evoked firing rates were quantified using generalized linear mixed-effects models (GLMMs) with spike count as the dependent variable with a Poisson distribution. The model specification `was n_events ∼ area*spike*rearing + offset(duration) + (1|unit) + (1|motif), where area*spike*rearing` defines fixed effects for brain region, cell type, rearing condition, and their interactions; offset(duration) defines a constant offset term for the duration of the interval in which the spikes were counted; and (`1|unit`) and (`1|motif`) define random intercepts for each unit and stimulus (omitted from the model for spontaneous rate). Contrasts, effects, and confidence intervals were calculated with emmeans. Contrasts are reported on the log ratio (LR) scale.

Signal and noise correlations were also computed using the number of spikes evoked in the interval between the start of each motif and 0.5 s after its end. Following Jeanne et al. (Jeanne et al., 2013), for each pair of simultaneously recorded neurons, the signal correlation was calculated as the Pearson correlation coefficient between the trial-averaged firing rates to the ten songs. The noise correlation was calculated by taking the Pearson correlation coefficient across trials of the same song and then averaging over songs. The noise correlation was corrected by subtracting the shift-predictor correlation, which is calculated the same way as the noise correlation but with trials that were not recorded simultaneously (Ponce-Alvarez et al., 2013). Only BS–BS and NS–NS pairs were considered, due to the difficulty of interpreting BS–NS correlations. Only auditory units (see below) were considered, and pairs recorded on the same channel were excluded. The effect of rearing condition on signal correlations was estimated using linear regression in R, with area, cell type, rearing condition, and their interactions as predictors. The same model was used for noise correlations. To test whether the relationship between signal and noise correlations was affected by rearing condition, we used a linear model with noise correlation as the dependent variable and area, cell type, rearing condition, signal correlation, and their interactions as fixed effects. Contrasts for CR versus PR were computed for each area and cell type using emmeans.

Discriminability was calculated for each unit by computing the pairwise similarity between all of its song-evoked responses. Responses were truncated to the duration of the shortest stimuli to remove information encoded in the response length. One stimulus was excluded because it was much shorter than the others. We used the SPIKE-synchronization metric (Kreuz et al., 2015) implemented in the pyspike Python package (v0.7.0; http://mariomulansky.github.io/PySpike/), yielding a 90 × 90 matrix of similarity scores that were used to train a *k*-nearest neighbors classifier (scikit-learn v1.3.2). That is, each trial was classified by finding the *k* most similar trials (the neighbors) and assigning it the label that was the most common in that group. We used *k* = 9, but other values produced nearly identical results. This yielded a 9 × 9 confusion matrix with one dimension corresponding to the stimulus that was actually presented and the other to the label that was assigned to the trial. Each unit received a score for the proportion of trials that were correctly classified. We used a GLMM with the number of correctly classified trials as the binomial dependent variable to quantify the effect of rearing condition. The model specification in R was `cbind(n_correct, 90 - n_correct) ∼ area*spike*rearing + (1|unit)`. Contrasts were computed using emmeans and are reported on the log odds ratio (LOR) scale. Discriminability was also used to classify neurons as auditory: a neuron was deemed to be auditory if its average score for all the songs was more than 1.64 standard deviations (the one-sided 95% confidence level for a z-test) above the mean for 500 permutations in which the song labels were randomly shuffled.

Selectivity was calculated for each unit as 1 minus the proportion of songs that evoked a firing rate that was significantly greater than the spontaneous rate. We used a generalized linear model with spike count as the Poisson-distributed dependent variable to determine significance. For the rare neurons that did not spike at all during the spontaneous intervals (2.6%, *n* = 70/2705), a single spike was added to the spontaneous count to regularize the model estimates. To quantify the effect of rearing condition, we used a GLMM with the number of songs with rates above spontaneous as the binomial dependent variable. The model specification in R was `cbind(10 - n_responsive, n_responsive) ∼ area*spike*rearing + (1|unit)`. Note that defining responsive motifs as “failures” gives a selectivity score where 0 is a unit that responds to all the stimuli and 1 is a unit that responds to none. Only auditory units were included in this analysis. Contrasts are reported on the LOR scale.

Stimulus reconstruction from population responses was implemented using a linear decoder. The model is similar to the spectrotemporal receptive field, in which the expected firing rate of a single neuron at a given time point *t* is modeled as a linear function of the stimulus spectrogram immediately prior to *t*, but in the linear decoding model, the relationship is reversed, and the expected stimulus at time *t* is modeled as a linear function of the response that follows. Using a discrete time notation where *s_t_* is the stimulus in the time bin around *t*, and *r_t_* is the response of a single neuron in the same time bin, then the expected value of the stimulus is given by

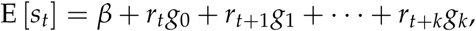

where *k* is the number of time bins one looks into the future, **g** = (*g*_0_, *g*_1_, …, *g_k_*) are the linear coefficients of the model, and *β* is a constant intercept term. If the errors are independent and normally distributed around the expectation with constant variance *σ*^2^, then this is an ordinary linear model. If there are *n* time bins in the stimulus, then the stimulus is a vector **s** = (*s*_0_, …, *s_n_*) drawn from a multivariate normal distribution. In vector notation,

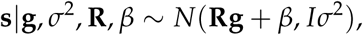

where **R** is the *n* × *k* Hankel matrix of the response. Without any loss of generality, the model can be expanded to include the responses of multiple neurons. If there are *p* neurons, then *r_t_*becomes a *p*-element vector (*r*_1,*t*_, …, *r_p_*_,*t*_), **R** becomes a *n* × *pk* matrix formed by concatenating the Hankel matrices for each of the neurons, and **g** becomes a *pk*-element vector.

In this study, the response matrix was constructed from the peristimulus time histograms (PSTHs) averaged over 10 trials using a bin size of 2.5 ms. We fit the model using all the PR neurons, all the CR neurons, or subsets of the PR or CR populations randomly selected without replacement. We used a *k* of 80, corresponding to lags from 0–200 ms. The response matrix was projected into a basis set consisting of 20 nonlinearly spaced raised cosines (Pillow et al., 2005). The width of each basis function increased with lag, giving higher temporal resolution at short lags and lower resolution at long lags. This allowed the inclusion of longer lags without exploding the number of parameters.

Stimuli were converted to time-frequency representations using a gammatone filter bank, implemented in the Python package gammatone (version 1.0) with 40 log-spaced frequency bands from 1–8.5 kHz, a window size of 5 ms, and a step size of 2.5 ms. Power was log-transformed.

We used ridge regression to estimate the parameters of the model because the number of parameters *pk* was typically larger than the number of time bins *n*. The decoder performance was estimated using a 10-fold cross-validation strategy. There were 10 data folds generated by holding out each of the songs. For each fold, 9 songs at the highest SNR (70 dB) were used to estimate *ĝ_λ_*, where *λ* denotes the ridge shrinkage penalty. We tested 30 *λ* values on a logarithmic scale from 10^−1^ − 10^7^. *ĝ_λ_* was used to decode the stimulus *S̃* from the responses to the held-out motif *R̃*, also at 70 dB SNR. The score for a given prediction was quantified as the adjusted coefficient of determination *R*^2^. The cross-validated prediction score was computed by averaging scores across folds for each value of *λ* and then taking the best score. To estimate noise invariance, we used the trained model to decode the stimulus from the responses to the held-out motif at the full range of SNR levels. Ridge regression and cross-validation were performed in Python with the scikit-learn library (v1.3.2).

## Results

### The zebra finch colony is a complex acoustical environment that masks individual songs

This study tested how the acoustical environment birds experience during development impacts auditory processing by comparing two rearing conditions. In the colony-reared (CR) condition, birds were housed with their families in individual cages situated in our breeding colony, where they were able to hear the songs and calls of many other individuals. In the pair-reared (PR) condition, birds were housed with their families in acoustic isolation chambers, where they could hear only the songs and calls of their parents and siblings. To characterize the differences between these environments, we collected acoustic recordings from one nestbox in each condition when juveniles were between 26–40 dph. The CR environment was much louder than the PR environment (Fig. 1A,D). On average, the median acoustic amplitude during the birds’ daytime hours was 50 ± 2 dB_A_ SPL (mean ± SD, *n* = 723 intervals of 10 min over 6 d) in the CR recording and 28 ± 5 dB_A_ SPL in the PR recording (*n* = 722). The average maximum amplitude was 88 ± 4 dB_A_ SPL in the CR recording and 77 ± 7 dB_A_ SPL in the PR recording.

**Figure 1.**
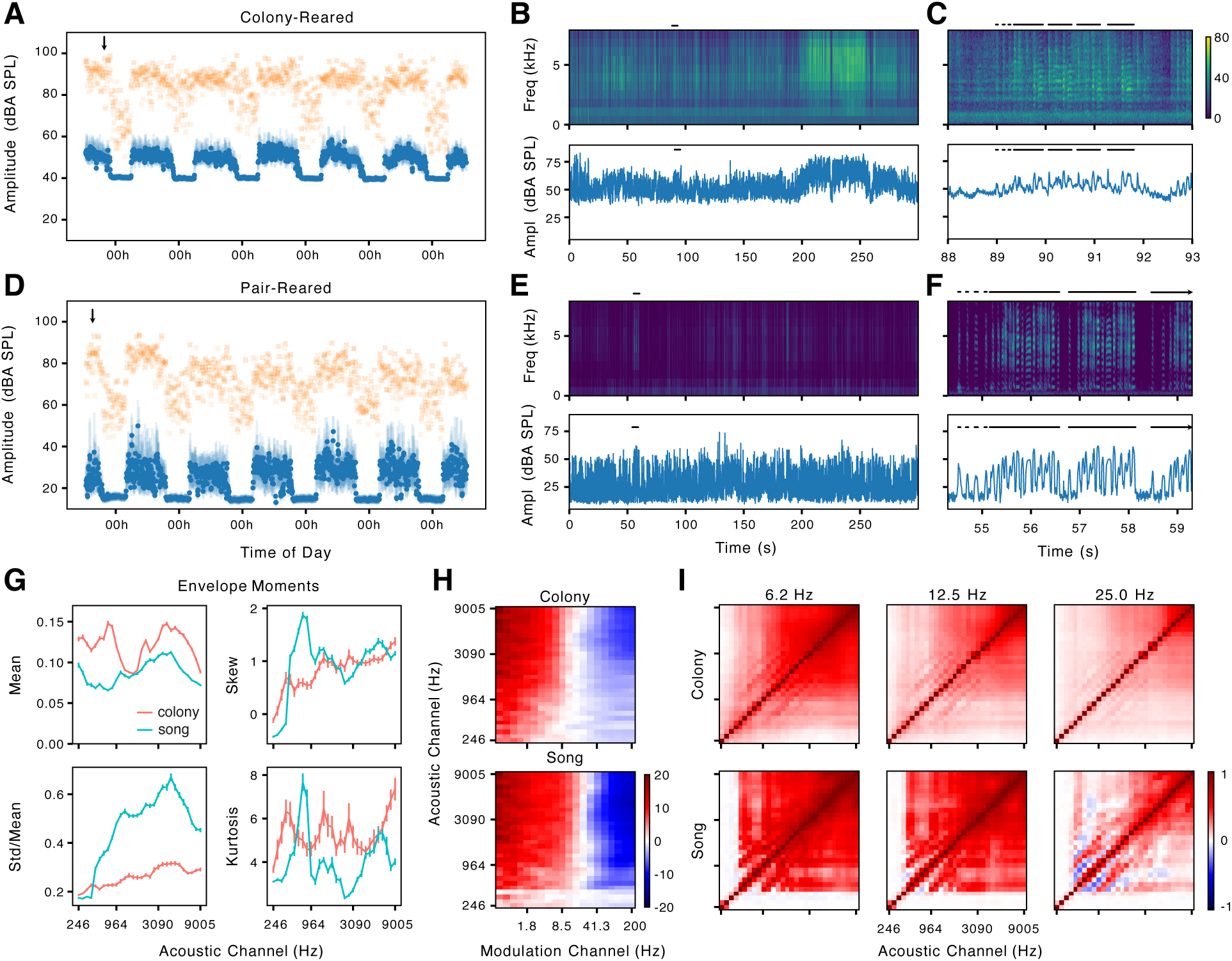
Statistics of acoustical environment for colony-reared and pair-reared chicks. (**A**) Sound pressure level (A-weighted) in 10-min intervals across several days in a recording from a nestbox in the colony. The clutch comprised four chicks that were 26–29 dph at the start of the recording, and the colony contained 70–72 animals (median 71). Blue dots, median; blue bars, interquartile range; yellow crosses, maximum. (**B**) Spectrogram (top) and amplitude (bottom) of colony noise during the interval marked by an arrow in A. The black bar marks a song by the father of this clutch. The loud sound between 200–250 s is a series of begging calls. (**C**) Detail of the interval marked by a bar in B. The bars mark introductory notes and syllables of the father’s song. (**D–F**) Same format as A–C but for a family housed in an acoustic isolation chamber. The clutch comprised six chicks that were 27–34 dph at the start of the recording. The color scale (dB SPL) is the same for all the spectrograms in the figure. The arrow in F indicates that the song continues beyond the displayed time period. (**G**) Marginal moments of the amplitude envelopes for log-spaced acoustic channels corresponding to locations on the basilar membrane. Lines show mean ± SE for samples from the colony recording in A–C (salmon; *n* = 66) and for samples from individual zebra finch songs (cyan; *n* = 62). (**H**) Modulation power in each acoustic channel, normalized by the variance of the envelope, expressed in dB relative to the same statistic for pink noise. (**I**) Correlations between the envelopes of the acoustic channels at different modulation rates.

The CR environment was also more complex than the PR environment. There was a nearly continuous background of sounds from birds and artificial sources in the colony (Fig. 1B,C). The frequencies below 1 kHz were dominated by wide-band, incoherent noise that sounded like it was produced by the room’s ventilation system. The higher frequencies also included some incoherent noise but were dominated by overlapping vocalizations. These were easily identified as zebra finch songs and calls by their characteristic harmonic structure, but it was rarely possible to distinguish specific types of vocalizations or distinct individuals. Even the song of the father, who was never more than 60 cm from the microphone, was almost impossible to identify except by having an experienced observer listen to the recording, because the spectrographic features of the song were heavily obscured by noise, and the amplitude modulations were only 10–15 dB above the noise floor (Fig. 1C). In contrast, the PR environment had long periods of silence during which the background amplitude was as low as 20 dB_A_ SPL. It was trivial to identify songs and calls, and the spectrographic features of songs were clear and distinct (Fig. 1E,F).

Thus, although a bird raised in the colony is exposed to a larger number and variety of conspecific vocalizations, at the same time this complex background is likely to mask the features of individual songs, including the song of a male juvenile’s tutor. Masking can occur not only at the level of the periphery (i.e., energetic masking) but at higher levels of the auditory system as well, when the background has similar higher-order spectrotemporal structure (i.e., informational masking (Kidd et al., 2008)). To characterize these higher-order features, we computed a series of statistics developed by McDermott and Simoncelli (2011) to characterize and synthesize sound textures. The statistics are based on a model of the auditory periphery that decomposes sound into logarithmically spaced frequency bands corresponding to locations along the cochlea (or basilar papilla, in the case of birds), followed by filtering of the amplitude envelope for each band, which reflects the tuning of more central auditory neurons to different modulation rates (Woolley et al., 2005). As seen in Figure 1G–I, the statistics of these higher-order features in our colony were broadly similar to the statistics for clean recordings of individual zebra finch song. In the colony, the envelopes for the acoustic frequency bands around 850 and 3500 Hz had the largest means, corresponding to the low-frequency noise and the higher-frequency vocalizations (Fig. 1B,C). The standard deviation of the envelopes was relatively low, but greater for higher frequencies, indicating that the amplitude of the vocalizations varied more over time than the incoherent noise. The skew and kurtosis were high across almost all of the frequency bands, suggesting that the recording had rare periods of high amplitude, perhaps corresponding to the vocalizations of individual animals. By comparison, the clean song recordings had less power at low frequencies, consistent with the absence of ventilation noise. The variance in frequencies spanning from about 500 to 7000 Hz was larger than in the colony recording, consistent with the lower noise floor in the sound isolation chamber. Skew and kurtosis were comparable to the colony recording, but there were large peaks around 850 Hz. Both colony noise and song had slow temporal modulations across a broad range of acoustic frequencies (Fig. 1H) and high correlations between acoustic frequency bands over a wide range of modulation rates (Fig. 1I), consistent with the broadband character of zebra finch song. There was a banding pattern in the cross-channel correlations for both colony noise and song that reflects harmonic structure.

### Colony-reared birds are better at recognizing conspecific song motifs embedded in colony noise

We examined the effects of the acoustical environment on auditory perception by training birds to discriminate between conspecific songs and then testing their recognition accuracy when these songs were masked by synthetic colony noise (Fig. 2A). Four CR and six PR birds of both sexes were first pre-trained on a two-alternative choice task to discriminate between two songs and then transferred to a novel set of four songs (Fig. 2B). Background noise was present in the training stimuli at an extremely low amplitude, 70 dB SNR. All of the animals in the study achieved a high level of discrimination performance (log odds ratio > 4.0; Fig. 2C,D). Training was followed by a pre-test phase in which birds were introduced to stimuli with a higher level of background noise (35 dB SNR), so that they would learn to respond to the foreground and ignore the background. The testing phase consisted of a series of blocks in which 80% of the stimuli were at 70 or 35 dB SNR and 20% were at a lower SNR, starting with 30 dB in the first block and decreasing in increments of –5 dB down to an SNR of –10 dB in the final block.

**Figure 2.**
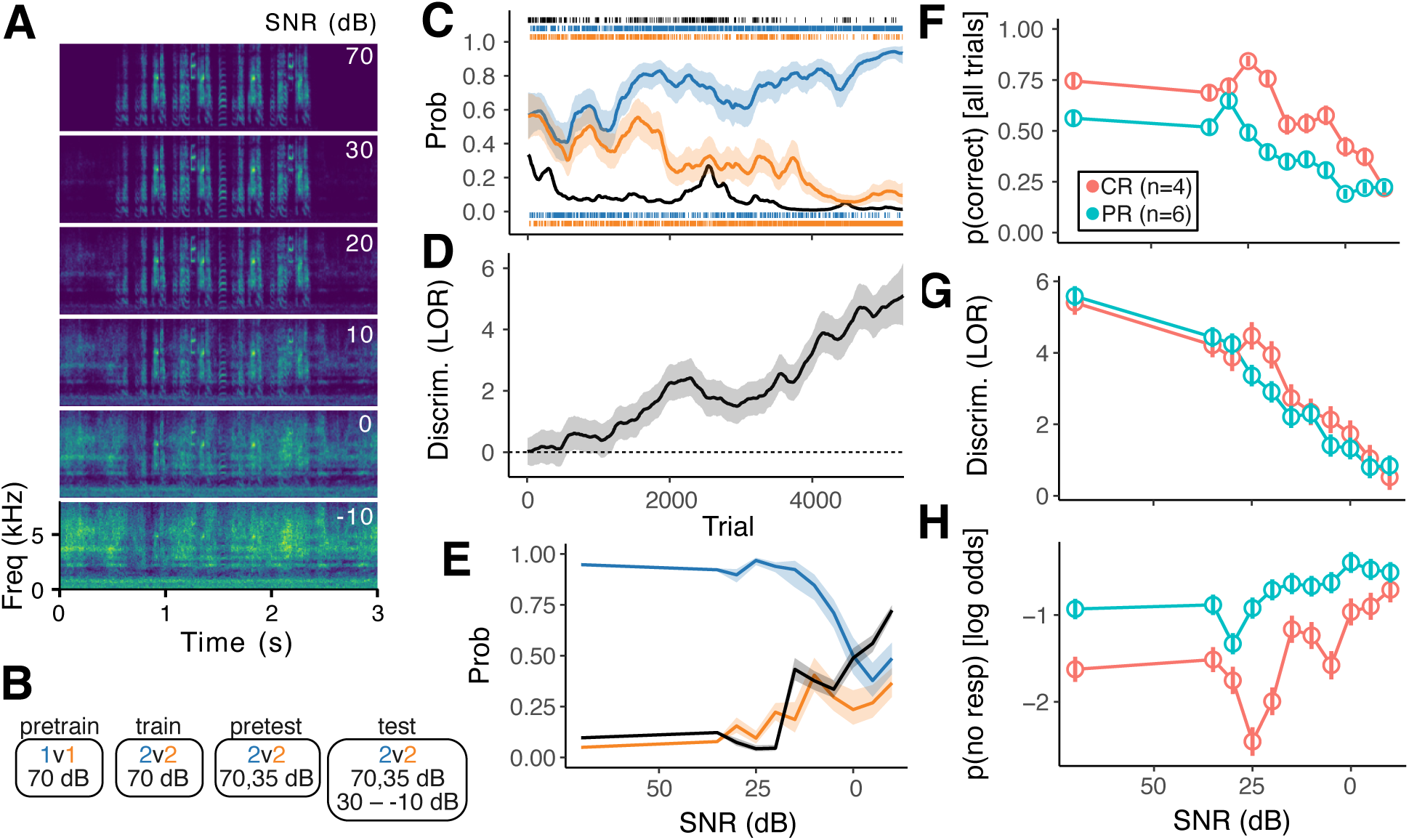
Operant training and recognition of songs embedded in colony noise. (**A**) Spectrograms of example stimuli used in the behavioral experiment. Stimuli comprised a single motif embedded in synthetic colony noise of varying amplitude. Each stimulus was presented in combination with five different samples of background noise. (**B**) Training and testing paradigm. Birds were trained to discriminate songs with inaudible background (70 dB SNR), trained to generalize to the 35 dB SNR condition, and then tested on songs with SNRs ranging from 30 to –10 dB SNR. (**C**) Learning curve for an exemplar CR bird during the training phase. Tick marks are individual trials. Black indicates trials where the bird did not respond, blue are trials where the stimulus was associated with the left key, and orange are trials with stimuli associated with the right key. The position of blue and orange ticks indicates whether the bird pecked left (top) or right (bottom). Blue, orange, and black lines indicate the maximum posterior probability of pecking left, pecking right, or not responding. Probabilities were estimated with a state-space model (see Methods). Shaded areas are 95% confidence intervals, not shown for p(no resp) for clarity. Correction trials are included in the trial count but are not shown or included in the analysis. (**D**) Posterior probability distribution of discrimination performance as the log of the odds ratio (odds of correctly pecking left on left-associated stimuli divided by the odds of pecking left on right-associated stimuli). Solid line shows maximum posterior probability, shaded areas the 95% confidence interval, estimated with state-space model. Dashed horizontal line, chance performance. (**E**) Response probabilities as a function of SNR during testing phase for the exemplar bird from (C). Lines indicate expected probability of pecking left (orange and blue) or not responding (black) with shaded regions indicating 95% confidence intervals, estimated with a generalized linear model. The probabilities of pecking left were only computed from the trials where the bird made a response. (F) Probability of making a correct response for PR and CR animals during testing phase, with failures to respond counted as incorrect. Symbols show estimated means (GLMM) and standard errors. Accuracy was higher for CR birds compared to PR birds (post-hoc *p <* 0.05) for all SNRs except 30 and –10 dB. Note that chance is not defined when counting non-responses. (**G**) Discrimination performance (log odds ratio) of PR and CR animals during testing phase. As in (C,D), discrimination is calculated only from trials in which the bird pecked one of the two keys. Symbols show estimated means (GLMM) and standard errors. Discrimination was greater for CR birds compared to PR birds (post-hoc *p <* 0.05) only at 25 and 20 dB SNR. (**H**) Log odds of PR and CR animals not responding during testing phase, same format as (F,G). PR birds were more likely to not respond (post-hoc *p <* 0.05) at all SNR levels except –10 dB.

As the SNR of the stimuli decreased, the animals were more likely to make mistakes, pecking left to songs that were associated with the right key, and vice versa (Fig. 2E). They were also increasingly likely to not respond on either key within 5 s after the end of the stimulus. Non- responses were punished the same as pecks to the incorrect key, so they were analyzed as incorrect responses. CR and PR birds both showed an overall decrease in accuracy as SNR decreased, but CR birds were more accurate than PR birds at all SNR levels except 30 and –10 dB (Fig. 2F). Averaging across all SNRs, the odds of a correct response were over twice as high in CR compared to PR birds (log odds ratio: 0.87 ± 0.24, *p* = 0.0003). This difference was almost entirely due to non-response rates. Excluding the non-response trials, CR and PR birds had similar discrimination performance except at 25 and 20 dB SNR, where CR birds were slightly but significantly better (Fig. 2G). CR and PR birds both showed the same trend of responding less as SNR decreased, but PR birds were less likely to respond than CR birds except at the lowest SNR of –10 dB (Fig. 2H). Averaging across SNRs, CR birds were half as likely to not respond as PR birds (log odds ratio: −0.71 ± 0.19, *p* = 0.0001).

### Birds raised in a complex acoustical environment have a more active auditory pallium

We recorded single-unit activity from the auditory pallium in a second cohort of birds comprising 9 CR and 7 PR adults of both sexes (4 CR males, 5 PR males). The stimuli comprised unfamiliar conspecific songs embedded in synthetic colony noise of varying amplitude. The avian auditory pallium consists of several quasi-laminar regions organized into columns with connectivity resembling the canonical neocortical circuit (Fig. 3A,B). Recordings were grouped into three broad subdivisions: the thalamorecipient areas L2a and L2b, the superficial areas L1 and CM (including medial and lateral), and the deep/secondary areas L3 and NCM. As has been previously reported (Meliza and Margoliash, 2012; Calabrese and Woolley, 2015), units in each of these regions had extracellular waveforms that were either narrow with larger peaks or broad with shallower peaks (Fig. 3C). Narrow-spiking (NS) cells are thought to be fast-spiking inhibitory interneurons, whereas broad-spiking (BS) cells are thought to be excitatory (Bottjer et al., 2019; Calabrese and Woolley, 2015). Table 1 shows the sample sizes from this experiment, including the number of cells of each type in each area. Auditory responses to song were diverse and complex (Fig. 3D), with different neurons responding to different songs with tightly time-locked peaks of activity or with slower modulations of firing rate. Across all three areas, BS neurons tended to have lower firing rates and to respond to fewer songs, whereas NS neurons tended to have higher firing rates and broader selectivity (Fig. 4A,B).

**Figure 3.**
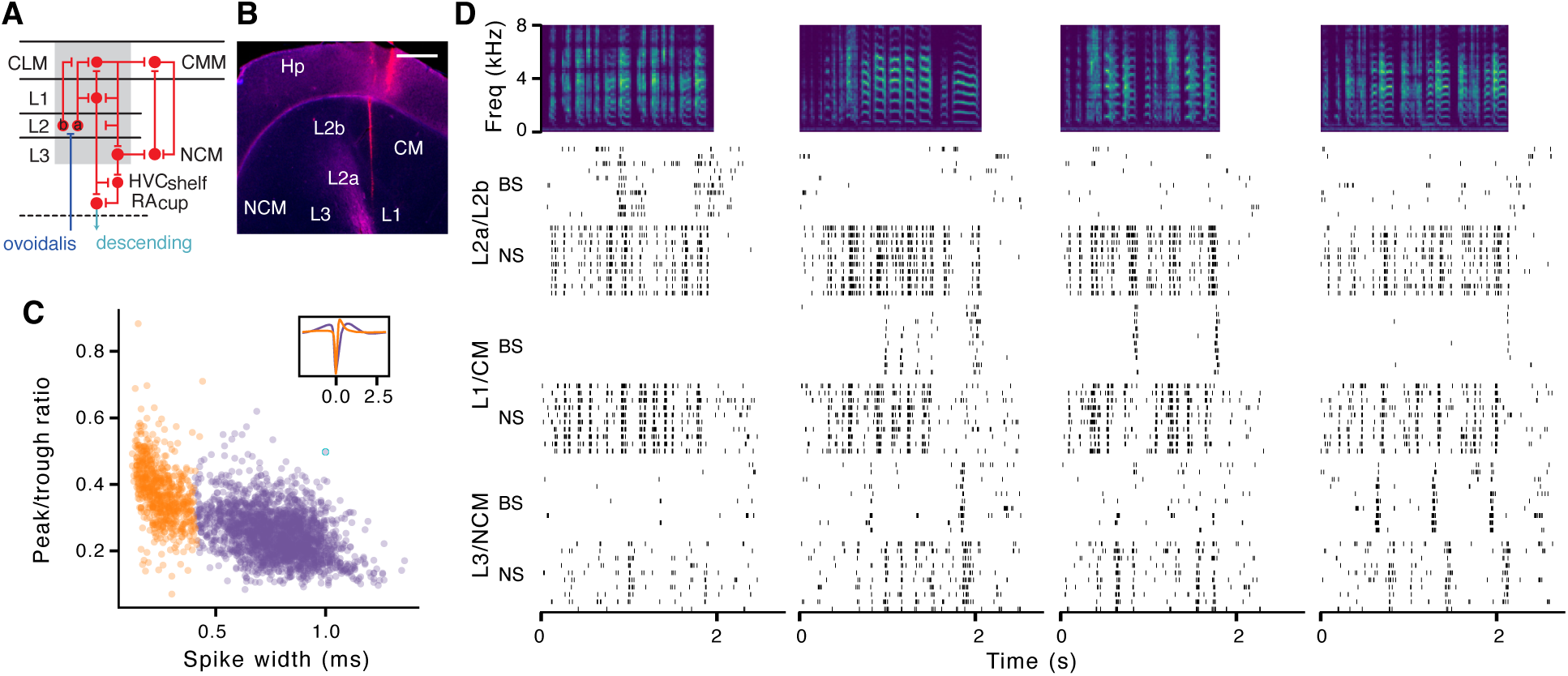
Example responses from zebra finch auditory pallium. (**A**) Architecture and connectivity of the auditory pallial areas examined in this study (adapted from Wang et al. 2010). Recordings were grouped into intermediate, thalamorecipient areas (L2a, L2b), superficial areas (L1, CLM, CMM), and deep/secondary output areas (L3, NCM). (**B**) Example histological image used to reconstruct recording location. The red track is DiI marking the location of one of the silicon electrode shanks; the other shank is in a different section. The section was counter-stained with DAPI (blue), but the signal was poor in this example. Electrodes were angled throughout the study as in the example so that all the recording sites were in L1 and CM, L2a and L2b, or L3 and NCM. Scale bar, 500 µm. (**C**) Peak-to-trough ratio and spike width for all units in the study. Units were classified as broad-spiking (BS, purple) or narrow-spiking (NS, orange). Inset shows average waveforms for the two classes. (**D**) Raster plots of example BS and NS units from each brain area responding to zebra finch songs (spectrograms at top).

**Figure 4.**
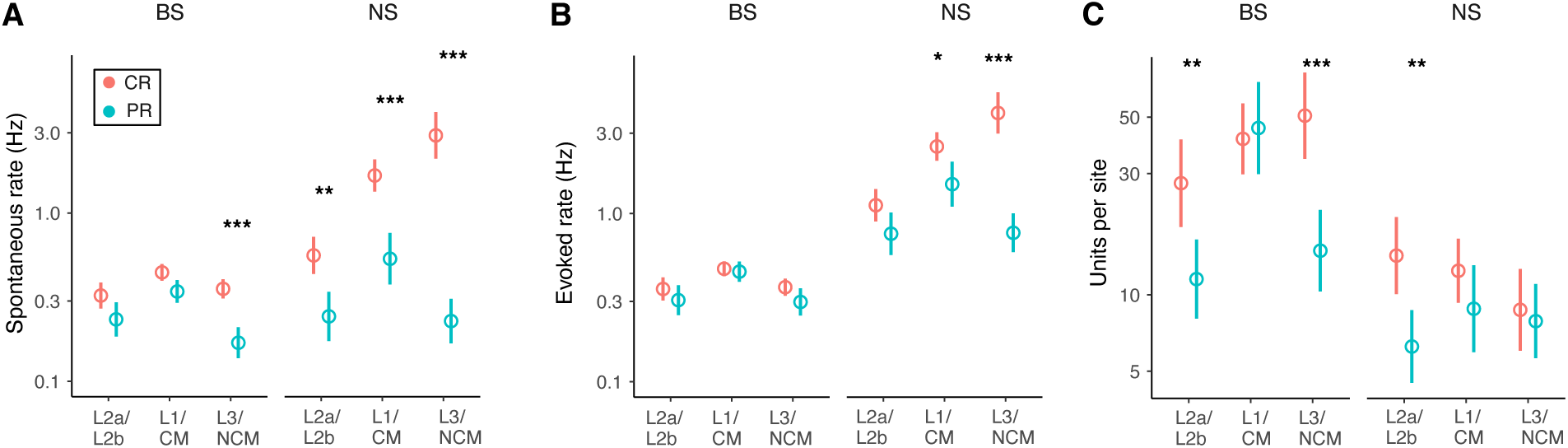
Effects of the acoustical environment on firing rates. (**A**) Average spontaneous firing rates of BS and NS neurons in each brain area for CR (salmon) and PR (cyan) adults. Circles indicate means; whiskers indicate 90% credible intervals for the mean (GLMM; see Materials and Methods). Asterisks indicate whether there is a significant post-hoc difference between means for PR and CR units in each condition (*: *p <* 0.05; **: *p <* 0.01; ***: *p <* 0.001). BS cells in CR birds had higher spontaneous rates compared to PR birds in L1/CM (log ratio [LR] = 0.26 ± 0.11; *z* = 2.2; *p* = 0.025) and L3/NCM (LR = 0.74 ± 0.15; *z* = 4.9; *p <* 0.001). NS cells in CR birds had higher spontaneous rates in L2a/L2b (LR = 0.83 ± 0.26; *z* = 3.2; *p* = 0.0011), L1/CM (LR = 1.14 ± 0.25; *z* = 4.5; *p <* 0.001), and L3/NCM (LR = 2.5 ± 0.27; *z* = 9.5; *p <* 0.001). Sample sizes for number of birds, recording sites, and single units are given in Table 1. (**B**) Average evoked firing rates of BS and NS neurons in each brain area. NS neurons in CR birds had higher evoked rates compared to PR birds in L1/CM (LR = 0.52 ± 0.22; *z* = 2.3; *p* = 0.02) and L3/NCM (LR = 1.64 ± 0.23; *z* = 7.0; *p <* 0.001). There was no significant difference between PR and CR birds in the evoked rate of BS neurons (all *z <* 1.53; *p >* 0.13). (**C**) Average number of of BS and NS neurons recorded per recording site in each brain area. CR birds had more BS units per site in L2a/b (LR = 0.87 ± 0.32; *z* = 2.6; *p* = 0.007) and L3/NCM (LR = 1.22 ± 0.33; *z* = 3.7; *p* = 0.002) and more NS units per site in L2a/b (LR = 0.82 ± 0.29; *z* = 2.8; *p* = 0.005).

Focusing our analysis first on the stimuli where the background was inaudible (70 dB SNR), we found a large effect of rearing condition on firing rates. Overall, CR neurons tended to have higher spontaneous and evoked activity than PR neurons, but the magnitude of the difference varied by area and cell type. The effect of rearing condition on spontaneous rate (Fig. 4A) was larger than it was on evoked rate (Fig. 4B), and the differences in spontaneous rate and evoked rate were largest for NS neurons in L3/NCM. We also found that we had recorded from fewer units in PR birds despite having a similar number of animals and recording sites (Fig. 4C; Table 1). This suggests that there were a large number of neurons in PR birds with firing rates below our threshold for inclusion (at least 100 spikes during the 58-minute recording session).

### Noise and signal correlations depend on rearing condition

The 3- to 10-fold greater firing rates of NS neurons in CR birds were not accompanied by a decrease in BS firing rates, as might be expected if the network connectivity were the same in both groups. This suggests that there were broader effects on the structure of inhibitory and excitatory networks. We tested this hypothesis by measuring the correlation of activity in pairs of simultaneously recorded neurons, with the expectation that increased inhibition in CR birds would decorrelate auditory responses, leading to sparser and more invariant representations of song. We calculated signal correlations, which measure how similarly tuned two neurons are, and noise correlations, which measure how much activity covaries when the stimulus is the same (Averbeck et al., 2006). Consistent with our hypothesis, we found that signal correlations were lower in CR birds compared to PR birds, although the magnitude and significance of the effect varied by brain region and cell type (Fig. 5A). BS neurons were less correlated in CR birds at earlier stations of the auditory hierarchy (L2a/L2b and L1/CM), whereas NS neurons were less correlated in CR birds at later stations (L1/CM and L3/NCM). We did not consider BS–NS pairs because of the difficulty of interpreting correlations between putatively excitatory and putatively inhibitory neurons.

**Figure 5.**
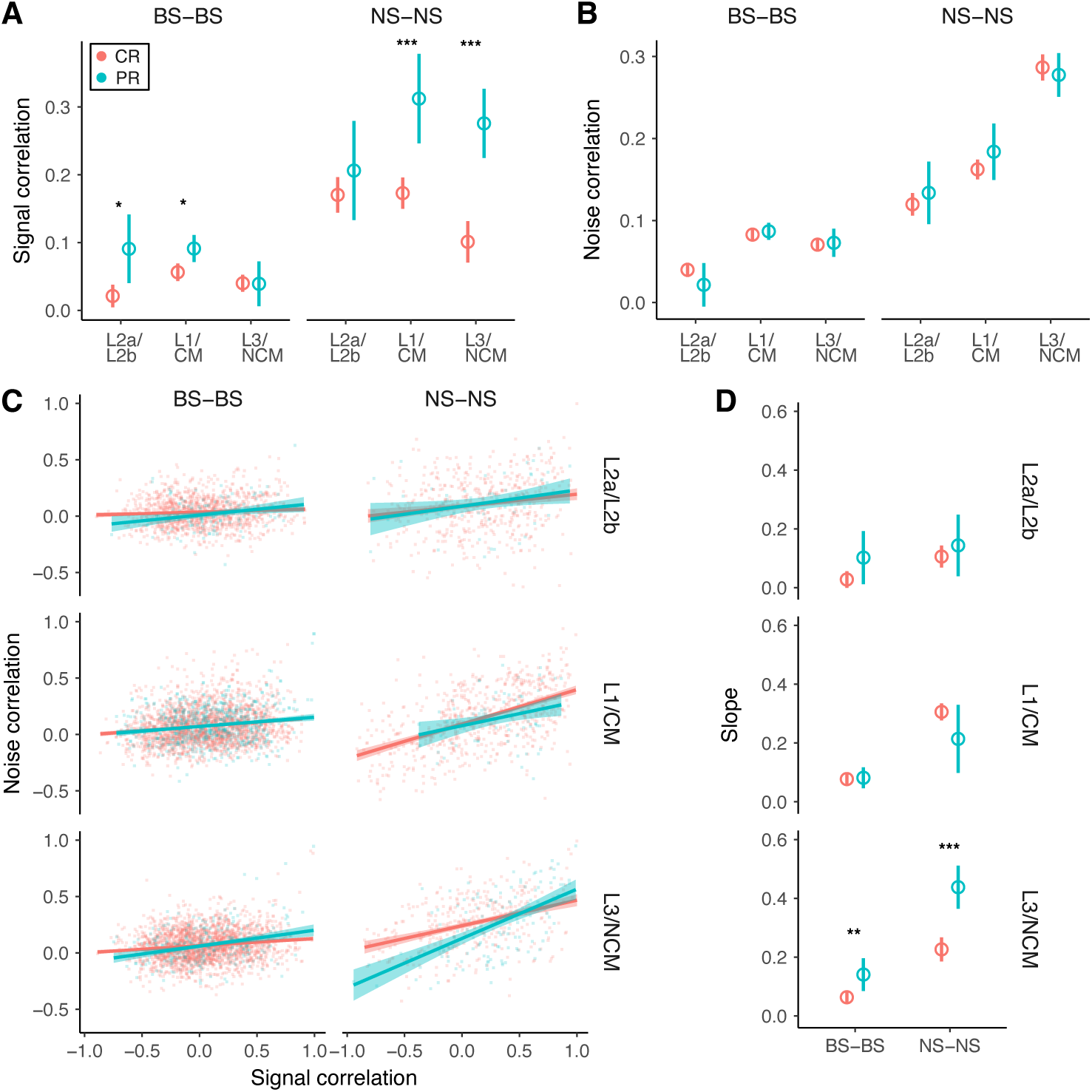
Pairwise correlations of auditory responses. (**A**) Signal correlations between pairs of BS neurons and pairs of NS neurons in each brain area for CR and PR birds. On average, BS neurons were less correlated with each other than NS neurons (*β* = −0.15 ± 0.01; *t*(8237) = −10.7; *p <* 0.001). Pairs of BS neurons in CR birds had lower signal correlations compared to PR birds in L2a/L2b (*β* = −0.07 ± 0.03; *t*(8237) = −2.1; *p* = 0.03) and L1/CM (*beta* = −0.03 ± 0.01; *t*(8237) = −2.4, *p* = 0.02). Pairs of NS neurons in CR birds had lower signal correlations compared to PR birds in L1/CM (*β* = −0.14 ± 0.04; *t*(8237) = −3.3; *p* = 0.001) and L3/NCM (*β* = 0.17 ± 0.03; *t*(8237) = −4.8; *p <* 0.001). (**B**) Noise correlations between pairs of BS neurons and pairs of NS neurons in each brain area for CR and PR birds. There was a significant interaction between area and cell type (*F*(2, 8237) = 34.8; *p <* 0.001), but none of the interactions involving rearing condition were significant (*F <* 0.8; *p >* 0.37). (**C**) Scatter plots of signal and noise correlations between pairs of BS neurons and pairs of NS neurons in each brain area for CR and PR birds. Colored lines are linear regressions with standard errors. (**D**) Regression slopes for C with 90% credible intervals. The slopes in CR birds are lower compared to PR birds in L3/NCM for both BS pairs (*β* = −0.08 ± 0.02; *t*(8225) = −2.5; *p* = 0.014) and NS pairs (*β* = −0.21 ± 0.04; *t*(8225) = −4.9, *p <* 0.001). All of the slopes were significantly greater than zero except for BS pairs from CR birds in L2a/L2b.

We did not observe a significant effect of acoustical environment on noise correlations (Fig. 5B), but there was a difference in the relationship between signal and noise correlations. As has been commonly observed in the cortex (Bair et al., 2001; Gu et al., 2011), noise correlations tended to increase with signal correlations in all three brain areas and both cell types (Fig. 5C,D). This relationship between noise and signal correlations is bad for coding efficiency, because similarly tuned neurons with correlated noise provide more redundant information and less reduction in uncertainty compared to neurons with independent or negatively correlated noise (Averbeck et al., 2006). In European starlings, learning to recognize the songs of other individuals inverts the typical relationship between signal and noise correlations, increasing the information ensembles of neurons carry about song identity (Jeanne et al., 2013). We observed a similar though smaller effect in L3/NCM (Fig. 5C,D), such that pairs of BS neurons and pairs of NS neurons with similar tuning (high signal correlations) had lower noise correlations in CR birds compared to PR birds.

### Auditory responses are more discriminable and selective in colony-reared birds

To characterize the effects of the acoustical environment on the functional properties of auditory neurons, we examined two complementary measures of sensory coding. The first measure, discriminability, assesses how reliably and distinctively a neuron responds to different stimuli. Neurons with more discriminable responses carry more information about which stimulus was presented. In this study, discriminability was quantified by using a timescale-free measure of spike train similarity (Kreuz et al., 2015) to make pairwise comparisons between the responses of the neuron in ten presentations of 9 songs (one song was excluded from this analysis because it was much shorter than the others). Examples of the similarity matrix that results from this comparison are shown for two example cells in Figure 6A. The similarity matrices were used with a k-nearest neighbors classi- fier to predict which stimulus was presented (Fig. 6B). Discriminability was taken as the proportion of trials that were correctly matched to the presented stimulus. The second measure, selectivity, assesses how narrowly a neuron is tuned. In this study we used a simple definition of selectivity based on the proportion of stimuli that evoked a response with a rate significantly greater than the neuron’s spontaneous firing (Fig. 6C). Selectivity and discriminability are complementary because they measure functional properties with an inherent tradeoff (Elie and Theunissen, 2015). As seen in the examples, highly selective neurons tend to have low discriminability scores, because a large proportion of stimuli evoke weak and inconsistent responses that are difficult to discriminate. Conversely, neurons that give distinctive responses to many different motifs will tend to have high discriminability scores but poor selectivity. Our results show evidence of this tradeoff at the level of cell types: averaging across areas and rearing conditions, BS neurons had lower discriminability compared to NS neurons (Fig. 6D) but greater selectivity (Fig. 6E).

**Figure 6.**
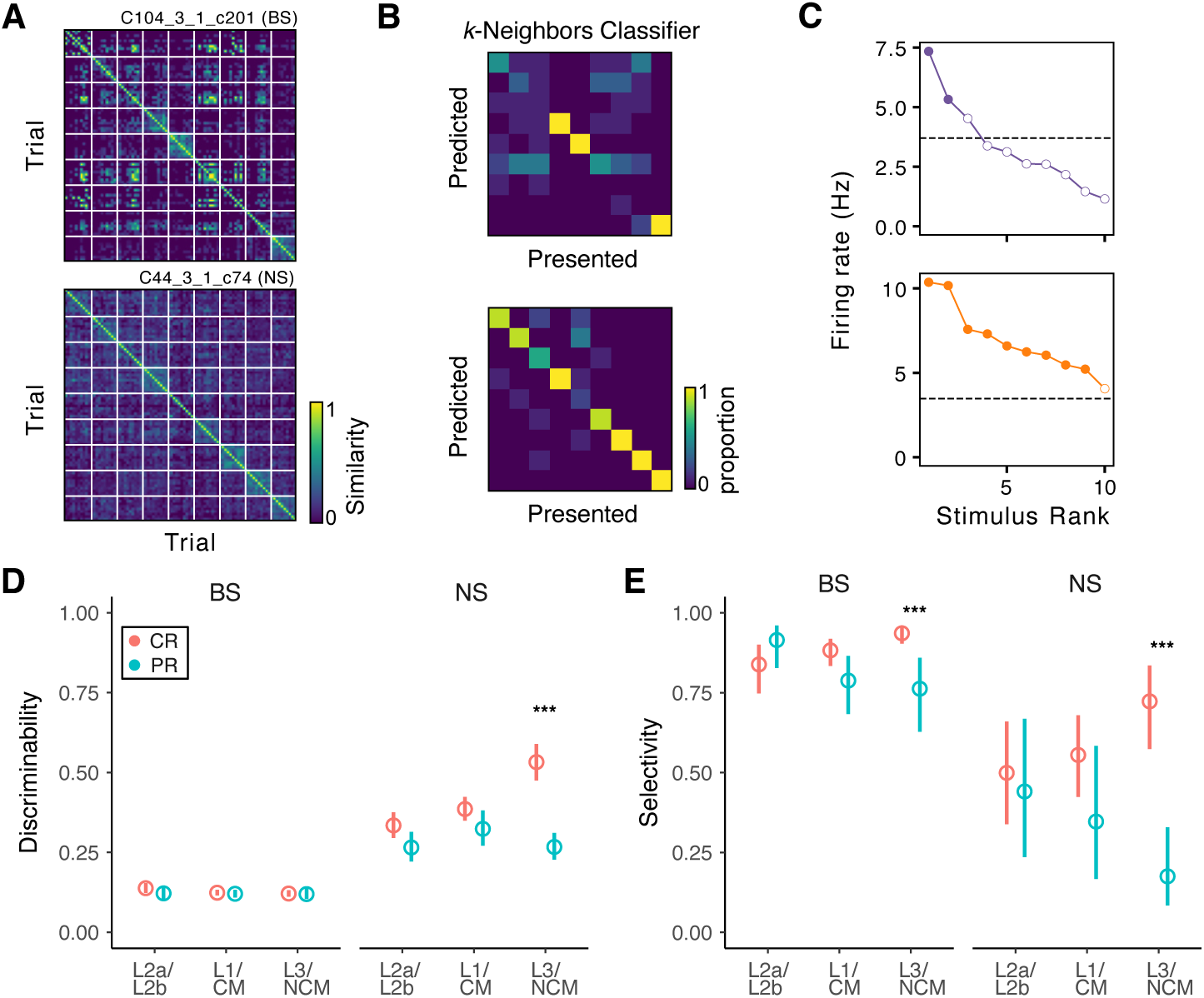
Discriminability and selectivity of auditory responses. (**A**) Spike-train similarity matrices between all pairs of trials for two exemplar units. The trials have been sorted by song, as indicated by the grid of white lines. Blocks along the diagonal correspond to comparisons between trials of the same song. (**B**) Confusion matrices from the predictions of a classifier model for each of the exemplars shown in A. Correct predictions correspond to the diagonal. The proportion of correct predictions was higher when spike train similarity was high between trials for the same song and low between trials for different songs. (**C**) Average firing rates of the two exemplar neurons in A–B evoked by each of the songs. The dashed line indicates the average spontaneous rate for that unit. Filled circles indicate responses significantly greater than the spontaneous rate (GLM; see Materials and Methods). (**D**) Average discriminability (proportion of trials correctly classified) for BS and NS neurons in PR and CR birds. The responses of BS neurons were less discriminable on average than NS responses (LOR = −1.3 ± 0.06; *z* = −21.5; *p <* 0.001). Discriminability of BS responses did not differ between PR and CR birds (LOR = 0.06 ± 0.06; *z* = 1.0; *p* = 0.31). Among NS cells, CR responses were more discriminable than PR responses in L3/NCM (LOR = 1.14 ± 0.19; *z* = 6.0; *p <* 0.001). (**E**) Average selectivity for BS and NS neurons in PR and CR birds. BS neurons were more selective on average than NS neurons (LOR = 2.0 ± 0.3; *z* = 8.4; *p <* 0.001). Selectivity was higher in CR birds compared to PR birds in L3/NCM for both BS (LOR = 1.5 ± 0.4; *z* = 3.2; *p* = 0.0014) and NS neurons (LOR = 2.5 ± 0.7; *z* = 3.9; *p <* 0.001).

Discriminability and selectivity tended to increase at later stages of the processing hierarchy, but only in colony-reared birds (Fig. 6D,E). This was observed as a statistically significant difference between CR and PR birds in L3/NCM, with CR birds having more discriminable NS neurons and more selective BS and NS neurons in this area.

### Stimuli are decodable from smaller numbers of neurons in colony-reared birds

We hypothesized that the less correlated and more discriminable responses in CR animals would lead to higher coding efficiency at the population level. To test this hypothesis, we measured how well the spectrotemporal structure of song stimuli could be decoded from neural activity in CR or PR birds. We used a simple decoder that was trained to predict the spectrum of the stimulus at a given time *t* as a linear function of the population response over the subsequent 200 ms (Fig. 7A). The decoder performance was quantified using 10-fold cross-validation, with 9 songs used for training and 1 song for testing in each fold. When trained with the entire set of CR neurons, the decoder was able to make a prediction of the spectrotemporal structure of the test stimulus at a high level of detail and accuracy (Fig. 7B). The predictions from the PR neurons were not as good, but there were also fewer PR neurons. To control for this confound, we tested decoder performance on randomly selected subsets ranging in size on a logarithmic scale from 40 to 927 units, with 100 replicates per size. Decoder performance indeed increased with population size, but subsets of the CR neurons outperformed equally sized subsets of PR neurons (Fig. 7C) except for the smallest population sizes (40 and 50 neurons). In terms of coding efficiency, smaller numbers of CR neurons were able to match the performance of larger numbers of PR neurons; for example, around 300 CR neurons carried as much information about the spectrotemporal structure of the stimulus as 927 PR neurons.

**Figure 7.**
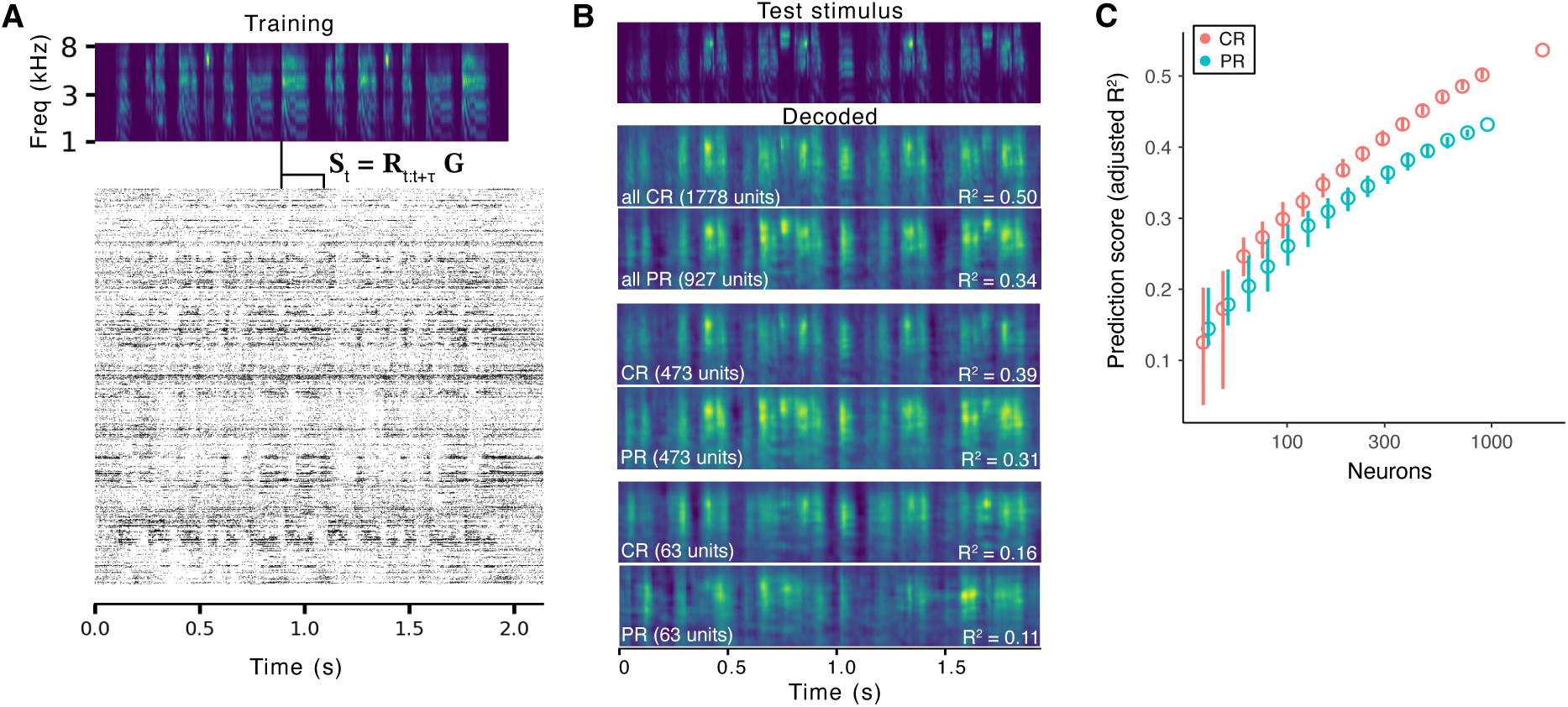
Decoding spectrographic features of song from population responses. (**A**) Raster plot of responses from all CR neurons to a zebra finch song. A linear decoder (*G*) was trained on 9/10 songs using ridge regression (see Methods) to predict the spectrum of the stimulus at each time point (*S_t_*) from the responses of the population in an interval immediately following that time (*R_t_*_:*t*+_*_τ_*). A gammatone filter bank was used for the spectrographic transform, resulting in a spectrum with log-spaced frequency bands that emphasizes lower frequencies. (**B**) To test decoder performance, the presented stimulus was decoded from the population response to the held-out song. Top spectrogram shows one test stimulus. Lower spectrograms are the decoder predictions for all CR units, all PR units, example large subsets of the CR and PR units, and example small subsets. *R*^2^ is the adjusted coefficient of determination for the prediction compared to the actual stimulus. The example subsets were chosen to match the median score for that size of population. (**C**) Decoder performance depends on the number of units used to train the model. The decoder was fit using subsets of the PR or CR units, sampled without replacement (*n* = 100 replicates per population size). Circles indicate median prediction score, whiskers the range between the 25% and 75% quantiles. Population sizes were matched across conditions, but the symbols are offset for clarity. The difference between median scores for CR and PR is significant for every population size (Wilcoxon rank-sum test, *p <* 0.01).

The PR population showed evidence of a sublinear relationship between population size and decoder performance, consistent with diminishing returns from combining neurons with similar tuning. This sublinear trend was also present for the CR units but less pronounced: the difference between CR and PR performance grew with population size, consistent with lower signal correlations and higher discriminability.

### Neural activity is less sensitive to a background of colony noise in colony-reared birds

CR birds were better than PR birds at recognizing songs masked by colony noise (Fig. 2), suggesting that the auditory pallium in CR birds is better able to filter out interference from background colony sounds. This effect could plausibly arise from the stronger and more precise inhibition in CR birds suppressing responses to background noises (see Discussion). To test this model, we examined how the auditory responses in CR and PR birds were affected by embedding stimuli in synthetic noise that matched the statistics of our zebra finch colony (Fig. 1). The foreground songs were presented in a shuffled sequence against a fixed background (Fig. 8A), so that each song was the foreground for 10 different segments of the background noise. As in the behavioral experiments, the amplitude of the background was varied to give SNRs between 70 and –10 dB. The effects of masking noise on individual units were diverse, but we often observed cells maintain the same firing patterns as the noise increased to around 5 dB SNR, after which the responses became weaker and more inconsistent (Fig. 8B).

**Figure 8.**
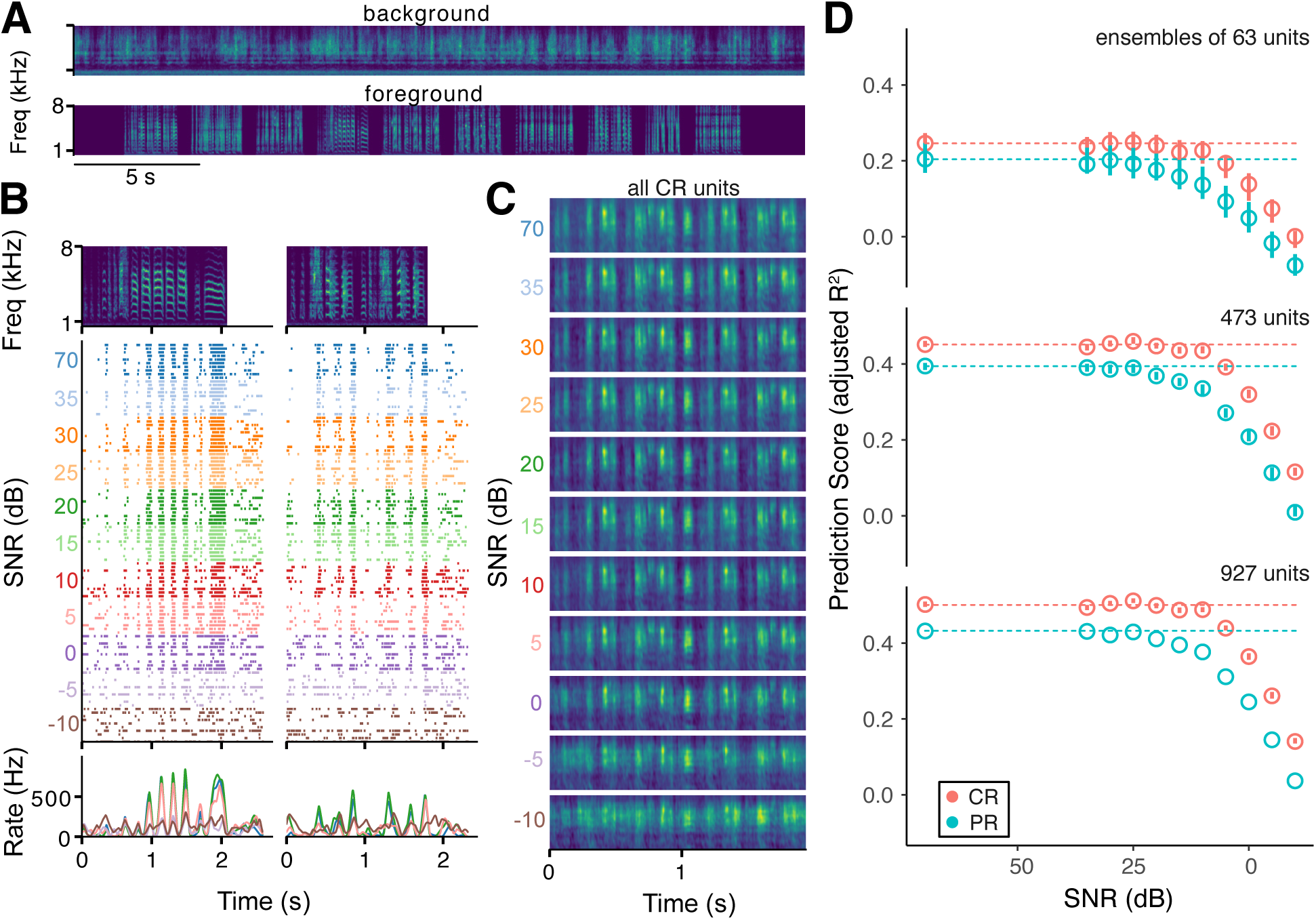
Decoding spectrographic features of song from population responses to song embedded in colony noise. (**A**) Spectrograms of background synthetic colony noise (top) and a representative sequence of 10 songs (bottom). The song order was shuffled so that each song was presented against a different segment of background in each trial at every noise level. (**B**) Raster plot of the responses of an exemplar unit to two songs (spectrograms, top) at different SNRs. Bottom plot shows smoothed firing rate histograms of the responses to selected SNRs. (**C**) Decoded spectrograms from the responses of all CR neurons at increasing noise levels. As in Figure 7, the decoder was trained on the responses of the population to the other nine songs at 70 dB SNR and then used to predict the stimulus from the responses to the tenth song embedded in noise at each SNR. (**D**) Decoder performance as a function of SNR for randomly selected subsets of the PR and CR units, sampled without replacement (*n* = 100 replicates per population size). Dashed lines indicate the median performance of the decoder on the 70 dB SNR responses. Whiskers indicate the range between the 25% and 75% quantiles. There are no error bars for the 927-unit PR ensemble, because this includes all the PR neurons.

To quantify sensitivity to noise at the population level, we used the same linear decoding approach as in the previous section. The decoder was trained on 9 songs presented without audible background noise (70 dB SNR) and then tested on responses to the remaining song at each SNR. Spectrograms decoded from responses to noise-embedded songs remained similar to the spectrograms decoded from responses to the noise-free songs up to about 0 dB SNR, at which point the decoded spectrograms began to take on characteristics of the background noise (Fig. 8C).

To compare CR and PR birds while controlling for differences in the number of recorded units, we used the same bootstrapping approach as in the previous section, with randomly selected subsets drawn without replacement from the CR or PR populations. The performance of decoders trained on CR units remained nearly the same up to about 5 dB SNR, whereas the PR-trained decoder predictions began to degrade at around 15 dB SNR (Fig. 8D). We observed this difference with all the population sizes we tested.

## Discussion

Species that communicate vocally require auditory systems that can reliably decode signals with complex acoustic features. In this study, we have shown that for a social songbird that breeds and raises its young in colonies, exposure to the social-acoustical environment of the colony during development profoundly affects how well birds can recognize conspecific songs in a behavioral task and how cortical-level neurons respond to these stimuli. The results are consistent with the idea that enriched experience is broadly beneficial to the development of sensory processing and support a model in which the developing brain adapts to the highly variable environment of many overlapping vocalizations by sharpening and strengthening inhibitory circuitry.

Compared to birds raised by pairs in acoustic isolation, colony-reared birds experienced a louder and more complex acoustical environment, with statistics that were similar to the statistics of song (Fig. 1). Thus, typical rearing conditions for zebra finches present a challenging “cocktail-party” problem for the developing auditory pallium to solve. The end result appears to be beneficial to the birds’ ability to recognize other individuals by their songs: CR and PR birds could both learn to discriminate novel conspecific songs in an operant task to a high level of performance, but CR birds were better at generalizing when songs were masked by synthetic colony noise at 20–25 dB SNR (Fig. 2G). More dramatically, CR birds were half as likely as PR birds to time out during the response window (Fig. 2H). This difference in non-response probability could reflect non-perceptual factors such as reduced cognitive control or increased aversion to risk (Baarendse et al., 2013), but it is also consistent with a difference in perceptual acuity, if PR birds are failing to respond when they are unable to clearly hear distinguishing features of the stimulus. Non-response probability increased in both groups as the SNR decreased, supporting the idea that birds are less likely to respond to more difficult stimuli. Further studies with other behavioral tasks would be needed to determine if colony-rearing also affects risk sensitivity or other non-perceptual processes.

Further supporting the conclusion that the social-acoustical environment of the colony affects the development of auditory perception, we observed large differences between CR and PR birds in the auditory pallium that were consistent with the behavioral observations. CR neurons had higher but less correlated firing rates, more discriminable firing patterns, more selective tuning, greater population-level coding efficiency for the spectrotemporal structure of song, and less sensitivity to background colony noise. We would expect all of these differences to positively contribute to perceptual discrimination.

The effects of colony-rearing were seen in multiple brain regions and cell types, but the largest and most consistent differences were in the narrow-spiking cells of NCM and L3. NCM and L3 are located next to each other in the auditory pallium, sharing an indistinct boundary (Fortune and Margoliash, 1992). They are both considered higher-order areas because neither receives direct input from the core of the auditory thalamus (Vates et al., 1996) and because neurons in both areas have nonlinear receptive fields and sharp tuning for small numbers of songs (Sen et al., 2001; Meliza and Margoliash, 2012; Calabrese and Woolley, 2015). L3 projects to NCM, which additionally exhibits experience- and learning-dependent responses consistent with a role in recognizing familiar individuals by their songs (Chew et al., 1996; Thompson and Gentner, 2010; Meliza and Margoliash, 2012; Schneider and Woolley, 2013) and in guiding sensorimotor learning for song production (Bolhuis et al., 2000; Stripling et al., 2001; Phan et al., 2006; Adret et al., 2012). Lesions to NCM disrupt discrimination of previously learned songs and recovery from noise-induced shifts in vocal production (but not new learning; Canopoli et al., 2014; Yu et al., 2023). Thus, experience-dependent changes to the functional properties of NCM and L3 neurons would be likely to affect perception and perceptual learning, and could be a major factor in the behavioral differences between CR and PR birds we observed here.

Although NS firing rates were higher on average in CR birds, so was their selectivity, and their firing patterns to different stimuli were more distinctive. Assuming that NS neurons are predominantly fast-spiking inhibitory interneurons (Calabrese and Woolley, 2015; Bottjer et al., 2019; Spool et al., 2020), this result indicates that exposure to a complex social-acoustical environment not only increases the overall inhibitory response to conspecific song but also sharpens its tuning for specific songs and the component features. Sharpened inhibitory tuning may explain why the evoked firing rates of BS cells were unaffected even as NS firing rates increased more than 3-fold (Fig. 4B). Indeed, we recorded from over three times as many BS units per NCM/L3 site in CR birds as we did in PR birds (Fig 4C). Because extracellular recordings are biased for units that spike often enough to be isolated during spike sorting, the most likely explanation for this large difference in the number of units is that there were many cells in the PR birds with extremely low firing rates. This is the opposite of what we would expect to see if increased inhibitory activity in CR birds led to a general dampening of responses, providing further evidence that exposure to the colony has fine-grained, specific effects on auditory processing. Our results strongly suggest that these effects are related to the reconfiguration of functional connectivity for greater coding efficiency. NS neurons in NCM/L3 had lower signal correlations with each other, which is suggestive of mutual inhibition. We also observed lower signal correlations in CR birds among BS neurons, but only in earlier stations of the hierarchy (L2a/L2b and L1/CM), suggesting that the acoustical environment may affect functional connectivity throughout the pallium and perhaps even at lower levels like the midbrain (Woolley et al., 2010). For both BS and NS neurons, the relationship between signal and noise correlations was altered, such that noise correlations were lower in similarly tuned pairs. This effect, which enhances coding efficiency (Averbeck et al., 2006), has also been reported in European starlings for stimuli that birds had learned to discriminate in an operant task (Jeanne et al., 2013). Confirming a link between functional connectivity and coding efficiency, we found that randomly selected populations of CR neurons carried more information about the spectrotemporal structure of song than PR populations of the same size (Fig. 7).

To synthesize these results, we propose a conceptual model based on mutual inhibition of excitatory ensembles (Harris and Mrsic-Flogel, 2013). NCM neurons receive excitatory inputs from upstream areas that are tuned to spectrotemporal features found in the songs of many individuals (Woolley et al., 2009; Meliza and Margoliash, 2012; Calabrese and Woolley, 2015; Kozlov and Gentner, 2016). NCM neurons are sensitive to the temporal sequencing of these features (Schneider and Woolley, 2013), which could result from the formation of Hebbian ensembles through repeated exposure to the unique songs of individual birds. Whereas CR birds are exposed to songs and calls from dozens of individuals, PR birds only hear the calls of their immediate family, the song of their father, and the (very similar) songs of their siblings. This impoverished experience would lead to less plasticity and the formation of fewer ensembles, explaining why there were so many fewer responsive neurons in PR birds. In contrast, the richer experience of CR birds would drive the formation of more ensembles and more robust excitatory connectivity. However, this rich experience comes with a cost: there is a nearly continuous background of overlapping vocalizations (Fig. 1), which would activate multiple ensembles simultaneously. Interference between ensembles would also result because songs share similar notes and syllables (e.g., the short harmonic stacks seen in all but one of the exemplar songs in Fig. 3D). One mechanism for achieving pattern separation with correlated noise and overlapping features is mutual inhibition, which creates winner-take-all dynamics that suppress activity from features in the background (Espinoza et al., 2018). In support of this model, blocking inhibition in NCM unmasks peaks of excitation in a context- and learning-dependent manner (Thompson et al., 2013; Schneider and Woolley, 2013). The experience-dependent increases in NS firing rates, selectivity, and discriminability we observed in this study are all consistent with stronger drive from excitatory ensembles onto inhibitory neurons, and the reductions in signal correlations among NS neurons and noise correlations among similarly tuned BS and NS neurons are consistent with a pattern of mutual inhibition. Furthermore, stronger mutual inhibition should also make responses more invariant to a background of colony noise. Our decoding analysis and our behavioral results confirmed this prediction: when decoding responses to noise-masked stimuli, CR neurons better preserved the foreground stimulus features compared to PR neurons (Fig. 8C,D), and CR birds were better at recognizing previously learned songs when they were masked by colony noise (Fig. 2F,G).

The results of this study are consistent with an extensive literature showing how the statistics of sensory experience shape postnatal development of the functional organization of the auditory cortex (Zhang et al., 2001; de Villers-Sidani et al., 2007; Zhou and Merzenich, 2008; Amin et al., 2013; Homma et al., 2020). With a few exceptions (Bao et al., 2013; Moore and Woolley, 2019), most of the prior work employed artificial stimuli, so the current study represents an advance in understanding how principles of experience-dependent plasticity apply to natural development in social, vocal communicators. However, a limitation of the experimental design is that there are multiple differences between the CR and PR conditions that are difficult to dissociate. Most notably, CR birds experience more signal and more noise: they hear many more songs than PR birds, but in an acoustical environment that masks the features of individual songs (Fig. 1). The noise in this study included mechanical sounds from the ventilation system in the colony and the vocalizations of other animals; we expect that the vocal noise makes a larger contribution to the effects we report because it is more likely to mask informative features of zebra finch song (Kidd et al., 2008), but developmental exposure to non-biological noises can affect auditory tuning as well (Amin et al., 2013; Homma et al., 2020). CR birds also have the opportunity for vocal and visual interactions with individuals outside their family, and there may be more diffuse effects of social density on brain development (Møller, 2010). It also remains to be seen to what extent the effects we observed are the result of an early sensitive period (Schroeder and Remage-Healey, 2021) or more recent plasticity (Yang and Vicario, 2015), and if they can be reversed (Furest Cataldo et al., 2023). Future work could begin to address these questions by controlling the timing and statistics of the acoustical background presented via loudspeaker to PR clutches.

Zebra finches have been studied extensively for their ability to imitate the song of a tutor heard during a critical period spanning approximately 25–65 days post hatch (Gobes et al., 2019). To control what songs they can imitate, birds in these studies are often isolated acoustically from the colony with their parents like our PR birds or with only female adults (e.g., Bolhuis et al., 2000; Phan et al., 2006). Males raised in such conditions produce accurate imitations of the tutor song (Tchernichovski and Nottebohm, 1998) despite, as our results indicate, having significant impairments in auditory processing and recognition behavior. This is perhaps unsurprising when one considers how important singing is to successful reproduction. Our sample size in this study was too small to examine potential effects of sex, number of siblings, and tutor song complexity, but another intriguing direction for future study is to examine whether birds hearing themselves and their siblings practicing during the later part of development moderates the effect of isolation from the larger colony.

Humans are social animals, but the range of social and acoustical environments they experience in infancy is enormous. Our results demonstrate that a complex environment has an impact on the development of auditory perception and the functional properties and connectivity of a higher-order auditory area. This finding aligns with the well-known link between an infant’s language experience and phonetic processing (Werker and Lalonde, 1988; Kuhl et al., 1992), while suggesting that there may be a broader dependence on the diversity and complexity of the acoustical environment in which this experience occurs. Understanding the mechanisms of this plasticity and its effects on perception and auditory learning may shed light on a broad range of language-learning impairments that are associated with deficits in auditory processing (Mody et al., 1997; Tsao et al., 2004; Ziegler et al., 2005; Pennington and Bishop, 2009).

## Acknowledgements

This work was supported by National Institutes of Health grant 1R01- DC018621, National Science Foundation grant IOS-1942480, and the University of Virginia Brain Institute. We would like to thank Jonah Weissman and Christof Fehrman for assistance with coding an early version of the linear decoder. Bao Le, Ayush Sagar, and Crystal Gong assisted in developing and testing the operant apparatus used in this study.

## Author Contributions

SMM and CDM conceived and planned the experiments. SMM carried out the experiments. SMM and CDM analyzed the data and prepared the figures. SMM and CDM wrote the manuscript.

